# Diagnostic images for mild cognitive impairment reveal biomarker status and abnormal scene processing

**DOI:** 10.1101/2023.11.30.569265

**Authors:** Yuetong Bai, Oliver Peters, Silka Dawn Freiesleben, Friederike Fenski, Josef Priller, Eike Jakob Spruth, Anja Schneider, Klaus Fliessbach, Jens Wiltfang, Claudia Bartels, Ayda Rostamzadeh, Wenzel Glanz, Stefan Teipel, Ingo Kilimann, Christoph Laske, Matthias H. Munk, Annika Spottke, Nina Roy-Kluth, Frederic Brosseron, Michael Wagner, Ingo Frommann, Falk Lüsebrink, Alfredo Ramirez, Peter Dechent, Stefan Hetzer, Klaus Scheffler, Renat Yakupov, Frank Jessen, Emrah Düzel, Wilma A. Bainbridge

## Abstract

Research on the impairment of episodic memory in Alzheimer’s disease often focuses on the processes of memory rather than the content of the specific images being remembered. We recently showed that patients with mild cognitive impairment (MCI), Stage 3 of Alzheimer’s disease, can memorize certain images quite well, suggesting that episodic memory is not uniformly impaired. Certain images, on the other hand, could not be memorized by MCI patients and were instead diagnostic for distinguishing MCI from healthy older adults. In this study, we investigate whether poor memory for diagnostic images is related to impaired neural processing in specific brain regions due to Alzheimer’s biomarker pathology. 64 healthy controls and 48 MCI participants in the DELCODE dataset performed a visual scene memory task during fMRI, with CSF Alzheimer’s disease biomarker data collected (i.e., amyloid and tau biomarkers). We found that diagnostic images have larger behavior-biomarker correlations for total tau, phospho-tau, Aβ42/Aβ40, Aβ42/phospho-tau compared to non-diagnostic images, suggesting that memory for these specific images are more affected by Alzheimer’s disease pathology. The fMRI data revealed an interaction effect between group membership (healthy control / MCI) and image diagnosticity (diagnostic / non-diagnostic scene images), with MCI participants having higher activation in scene processing regions (parahippocampal place area, retrosplenial cortex and occipital place area) for diagnostic images than non-diagnostic images. In contrast, healthy controls showed no differences in processing between diagnostic and non-diagnostic images. These results suggest that MCI individuals may engage in inefficiently heightened encoding activation for these diagnostic images. Our results show that special “diagnostic” images exist that can reveal amyloid and tau pathology and differences in neural activity in scene regions.

## Introduction

Memory loss is the earliest major symptom in the progression of Alzheimer’s Disease. Before developing into dementia, many individuals first experience amnestic mild cognitive impairment (MCI), where their memory performance falls significantly below their age-norm. While we know that individuals with MCI generally have worse memory than cognitively normal controls, less is known about the mechanisms of this impairment. Does episodic memory decline equally for all memoranda or do certain memoranda suffer more than others? If the latter is true, which brain regions contribute to content-specific memory loss? Here, we addressed these questions with a content-based exploration of Alzheimer’s biomarker and functional neuroimaging data from healthy controls and MCI individuals.

Studies on episodic memory impairment in the face of neurodegeneration have primarily focused on memory processes and tasks rather than memory content (Grande et al., 2021). Typical tasks for memory assessment are designed and performed largely independent of the specific contents to be remembered (Costa et al., 2017). In other words, little effort has been made to evaluate whether specific stimuli are more effective than others in revealing clinical differences. In some cases, different participants are assigned with different stimuli for the same task and the performance is assumed to be comparable. However, this assumption may not be true, as memory for certain contents may be specifically damaged in some dementia stages.

Prior works have characterized how memory contents are differentially remembered by people by examining the *memorability* of images. Memorability is an intrinsic attribute of a visual stimulus that describes the likelihood of successful memory in the general population. For scene images (Isola et al., 2011a), face images (Bainbridge et al., 2013), words (Xie et al., 2020), and many other stimulus types, some items are consistently remembered better than others. Importantly, memorability is driven by characteristics of the stimuli themselves and remains stable across different people. This consistency of memorability is observed in various memory and perceptual task paradigms (Goetschalckx et al., 2017; Broers & Busch, 2020; Bainbridge, 2020) and across different presentation times and delay intervals (e.g., Borkin et al., 2015; Isola et al., 2013), and is separable from the surrounding image context (Kramer et al., 2023). Memorability is also independent of other cognitive attributes such as attentional state (Wakeland-Hart et al., 2022) and encoding strength or motivation (Bainbridge, 2020). Thus, memorability is an intrinsic attribute for the stimulus that is relatively robust across tasks, contexts and observers. In exploring what makes a stimulus memorable, studies have suggested that semantic attributes explain a large proportion of variance (Kramer et al., 2023), while visual attributes explain much less variance (Isola et al., 2013; Dubey, et al., 2015).

Neuroimaging studies have shown that image memorability may influence how specific items are processed and represented throughout visual perception and memory processes. Functional magnetic resonance imaging (fMRI) studies compared activity associated with viewing memorable versus forgettable images while controlling for other low-level visual features and semantic features (Bainbridge et al., 2017). Participants showed more activity in inferotemporal cortex (IT) and medial temporal lobe (MTL) for images that are memorable versus forgettable on average, even when they themselves did not successfully remember the memorable images. These regions engage in high-level visual perception and memory encoding, suggesting that memorability may touch upon both processes. Moreover, the memorability effect is distinct from low-level vision and individual-specific memory processes. Even in non-human primates, image memorability can be predicted from the population magnitude response of IT neurons (Jaegle et al., 2019). The current hypothesis on why the brain shows such sensitivity to memorability suggests that image memorability is a high-level visual property that reflects the prioritization of certain information for processing and memory (Xie et al., 2020).

While many studies have explored memorability in cognitively normal participants, researchers are only starting to test how memorability for the same contents may differ between healthy people and memory-impaired individuals. A recent study tested whether a difference in memorability scores for a stimulus exists across groups, and whether it can be used as a diagnostic tool for MCI assessment (Bainbridge et al., 2019). This study examined a specific set of images for which a large memorability difference was observed between HC and MCI participants. If neurodegeneration causes uniform loss of all memory contents, we should expect a general drop in memorability scores from HCs to MCI participants but find few images that show a drastic memorability difference between the two groups. That is, the images that HCs remember best should also be the images that MCI participants remember best. However, the researchers found that *different* images were memorable and forgettable in the HC and MCI groups, with a change in memorability patterns. While some images were similarly memorable or forgettable to both HCs and MCI participants, a set of images that was memorable to HCs became forgettable to MCI participants (Figure 1). While memory for other images remained relatively intact, MCI participants showed targeted difficulties in remembering these specific images, indicating that these images may tap into the most vulnerable components of episodic memory. Indeed, classification models based on a participant’s memory for these “diagnostic” images significantly predicted whether a participant belonged to a HC or MCI group better than classification models based on their memory for other images. Cognitive assessments designed to include more of these diagnostic images may have better sensitivity in detecting the subtle memory impairments in early stages of Alzheimer’s dementia.

**Figure 1.**
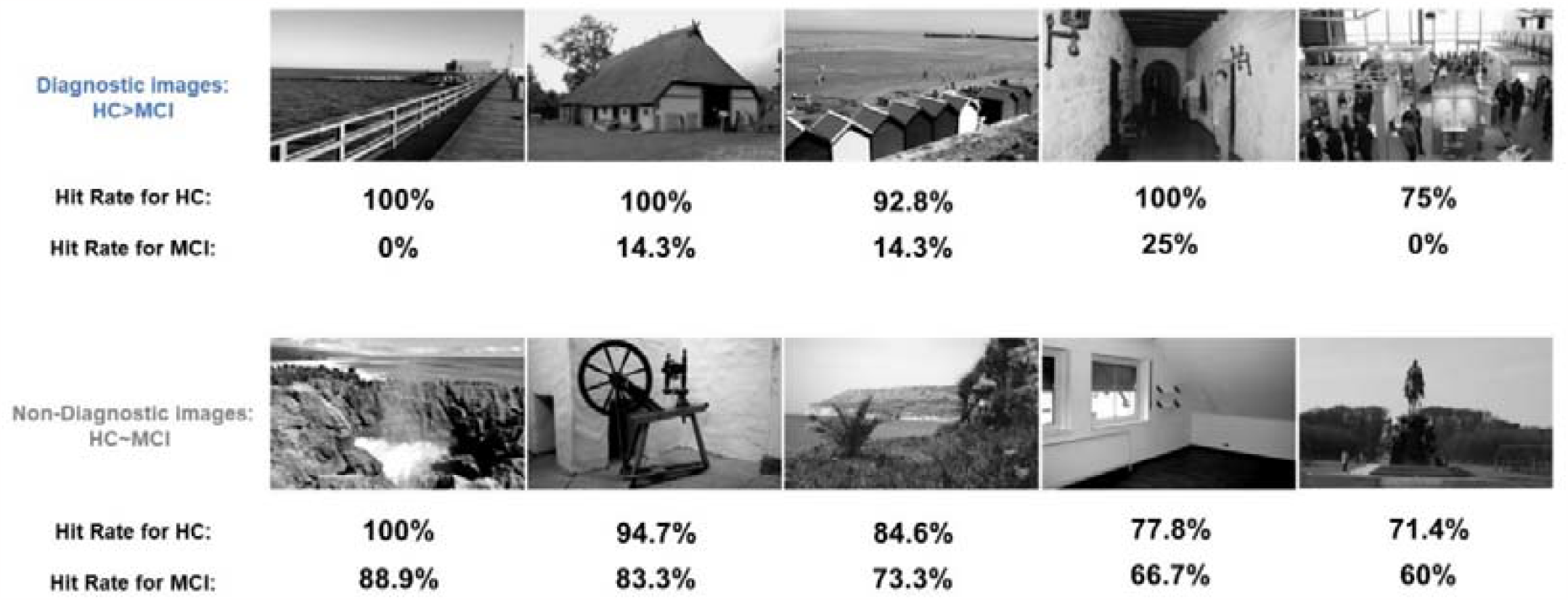
Sample diagnostic and non-diagnostic images in the current dataset. Diagnostic images (upper row) are images for which HCs have a much higher hit rate (HR) than MCI individuals. Non-diagnostic images (lower row) are images for which HCs have a slightly higher HR than MCI individuals, close to the average performance difference between HCs and MCI.

Although behavioral studies suggest that diagnostic images have the potential to improve clinical practices, little is known about how image diagnosticity relates to an individual’s Alzheimer’s Disease biomarker status or brain activity. Understanding this relationship could help to explain what specific brain pathologies are highlighted by diagnostic images. Amyloid and tau-pathology, hallmark biomarkers of Alzheimer’s disease, are commonly measured by cerebrospinal fluid (CSF) (e.g., total tau, phospho-tau and Aβ42, Aβ42/Aβ40, Aβ42/phospho-tau). Both pathologies cause neural dysfunction due to synaptic impairment and atrophy and cell loss in brain regions crucial for successful episodic memory (Jagust, 2018). As Alzheimer’s disease pathologies are revealed by biomarker and neuroimaging data, diagnostic images may show a stronger relationship with certain biomarker or functional activity patterns, which can explain why these images become more diagnostic.

In this study, we investigated whether poor memory for diagnostic images in MCI due to Alzheimer’s disease is related to impaired neural processing in specific brain regions due to Alzheimer’s biomarker pathology. We integrate behavioral, CSF biomarker and fMRI data to explore whether images with high memorability differences between HC and MCI participants (diagnostic images) could better indicate Alzheimer’s disease pathology. Using a cross-validation approach, we find that for several biomarkers, memory performance for diagnostic images showed significantly higher correlations with biomarker status than non-diagnostic images. We also find that for diagnostic images, MCI participants have higher activation in scene processing regions despite their worse memory performance, in contrast with HCs who show no activation differences between diagnostic and non-diagnostic images. These results suggest that diagnostic images could reveal Alzheimer’s disease pathologies better than other images and that they pose a particular neural processing challenge to scene regions.

## Results

### Consistency of hit rate and image diagnosticity in HC and MCI individuals

We analyzed the subset of participants from the DZNE Longitudinal Cognitive Impairment and Dementia Study (DELCODE) who had behavioral, biomarker, and fMRI data available (Table 1; Jessen et al., 2018). 64 HC and 48 MCI participants completed a scene memory task and had their CSF Alzheimer’s disease biomarker status measured. During the encoding phase (in scanner), they viewed novel scene images and completed indoor/outdoor judgements. In the recognition phase (out of scanner), they rated 1 to 5 on whether they have seen these images as well as some new images. From the memory performance of HC and MCI individuals, we identified sets of diagnostic and non-diagnostic images, where diagnostic images were those with the greatest performance differences between groups, while non-diagnostic images were those with the smallest differences (see Methods).

**Table 1.** Characteristics of participants. P-values denote results of independent-sample t-test comparisons between HC and MCI group. HC: healthy control group; MCI: mild cognitive impairment group; MMSE: Mini-Mental State Examination; Aβ42/40: ratio of Aβ42 to Aβ40; Aβ42/Phospho-tau: ratio of Aβ42 to phospho-tau

|  | HC n | MCI n | HC Mean(SD) | MCI Mean(SD) | P |
| --- | --- | --- | --- | --- | --- |
| Age | 64 | 48 | 68.70(4.97) | 72.33(5.00) | <0.001 |
| Years of education | 64 | 48 | 14.52(2.65) | 13.63(2.71) | 0.08 |
| MMSE | 64 | 48 | 29.38(0.85) | 27.94(1.56) | <0.001 |
| Total-tau | 64 | 48 | 348.53(141.7) | 538.60(261.1) | <0.001 |
| Phospho-tau | 64 | 48 | 46.98(15.12) | 67.18(29.43) | <0.001 |
| Aβ42 | 64 | 48 | 823.92(274.9) | 646.09(311.9) | <0.001 |
| Aβ42/40 | 64 | 48 | 0.097(0.02) | 0.077(0.03) | 0.002 |
| Aβ42/Phospho-tau | 64 | 48 | 18.26(4.92) | 12.20(7.81) | <0.001 |

In the recognition task, as expected from their differing pathology, HC participants outperformed MCI participants: HC participants on average had a HR of 75.28% (SD = 11.69%) while MCI participants had a hit rate of 63.63% (SD = 19.94%). We identified diagnostic and non-diagnostic images from the pool of 835 total stimulus images based on the behavioral performance of all participants. For the 100 diagnostic images, for which HC participants had much better performance than MCI participants, HC participants had an average HR of 89.27% (SD = 13.07%) and MCI participants had an average HR of 29.69% (SD = 15.34%), with an average hit rate difference between HC and MCI participants of 59.58% (SD = 14.34%). For the 100 non-diagnostic images identified from the whole sample, for which the HR difference between groups were close to the average of all images, HC participants had an average HR of 75.11% (SD = 15.34%) and MCI participants had an average HR of 62.45% (SD = 15.14%), with an average hit rate difference between HC and MCI participants of 12.66% (SD = 4.24%).

If certain intrinsic attributes of an image affect how it is remembered by healthy controls and participants, we should expect some level of consistency for the memory performance within HC and MCI groups, and for the performance difference between groups. That is, if an image is memorable/forgettable for one set of healthy controls, it should also be memorable/forgettable for another independent set of healthy controls (and this should also hold true for MCI participants). To examine this, we conducted a split-half consistency analysis with 1000 rounds of cross-validation. The average Z-transformed Spearman correlation coefficient between the two halves was ρ = 0.20 (SD = 0.05) for HC participants and ρ = 0.097 (SD = 0.07) for MCI participants, both significantly above 0, indicating a consistency in memory (one-sample t-test of Z-scored coefficients: p < 10^−10^). We also examined whether the performance differences between the two groups were consistent. The average Z-transformed Spearman correlation coefficient for diagnosticity was ρ = 0.10 (SD = 0.11) and was significantly higher than zero (p < 1^-10^). These results show that memory performance for a given image is highly consistent within the HC and MCI groups. Further, the difference in performance between HCs and MCIs for a given image is also consistent, demonstrating that images have an intrinsic “diagnosticity”.

### Overall prediction of memory performance based on biomarkers

We calculated the correlation between the five CSF biomarkers included in our analysis. As shown in Table 2, While Aβ42 doesn’t significantly correlate with total tau and phospho-tau, the other biomarkers strongly correlate with each other. We next examined the general power of CSF biomarker status to predict participants’ HR on all images, which will demonstrate how closely biomarker status is linked with task performance. Separate linear regression models were used to predict a participant’s HR based on each of the five target biomarkers. All biomarkers except for phospho-tau were significant predictors of hit rate (Aβ42: R^2^ = 0.023, p = 0.02; total tau: R^2^ = 0.035, p = 0.004; phospho-tau: R^2^ = 0.011, p = 0.11; Aβ42/40: R ^2^= 0.023, p = 0.02; Aβ42/phospho-tau: R^2^ = 0.026, p = 0.01). These results first confirm that there is a basic relationship between CSF biomarker and behavioral performance before we test this relationship separately for diagnostic and non-diagnostic images.

**Table 2.** Correlation between CSF biomarkers. t-tau: total tau; p-tau: phospho-tau; Aβ42/40: ratio of Aβ42 to Aβ40; Aβ42/p-tau: ratio of Aβ42 to phospho-tau **: p < 0.001

| | Aβ42 | t-tau | p-tau | Aβ42/40 | Aβ42/p-tau |
| --- | --- | --- | --- | --- | --- |
| Aβ42 | | | | | |
| t-tau | -0.13 |  |  |  |  |
| p-tau | -0.13 | 0.88** |  |  |  |
| Aβ42/40 | 0.76** | -0.56** | -0.58** | | |
| Aβ42/p-tau | 0.65** | -0.67** | -0.73** | 0.90** | |

### Diagnostic images provide a stronger link between behavior and biomarker status

The diagnostic images are characterized by a much better performance for HC and much worse performance for MCI than any other images. To verify that these behavioral differences really reflect impaired memory associated with neurodegeneration, we examined whether the behavioral performance of diagnostic images is more relevant to biomarker status indicative of MCI. Specifically, we compared the correlations of behavioral performance and biomarker status between diagnostic and non-diagnostic images (Figures 2 & 3). Importantly, we used a cross-validation approach, so we tested the ability to predict participants’ biomarker status based on diagnostic images determined from a separate set of participants. For four of the five biomarkers we examined, the diagnostic set had significantly larger correlations between behavior and biomarker measures than the non-diagnostic set (total tau: mean difference = -0.050, p < 0.001; phospho-tau: mean difference = -0.060, p < 0.001; Aβ42/40: mean difference = 0.029, p < 0.001; Aβ42/phospho-tau: mean difference = 0.040, p < 0.001). As the only exception, for Aβ42, the correlations from diagnostic images were not significantly different from non-diagnostic images (mean difference = -0.0008, p = 0.18). In conclusion, the behavioral performance calculated from the diagnostic images consistently showed a stronger link with CSF biomarker measurements compared to the non-diagnostic images for all biomarkers except for Aβ42. Further, these links were found with a separate set of participants than those used to measure image diagnosticity, demonstrating that these results are generalizable across people. As a whole, these results indicate that the diagnostic images are overall more sensitive for predicting the amyloid and tau pathology of both HC and MCI participants, which can be directly related to neurodegeneration specific to Alzheimer’s dementia. As these biomarker results serve as evidence that visual memory processes for the diagnostic images may be more impaired by neuronal pathology, we further examined how memory processes are impaired by analyzing fMRI data.

**Figure 2.**
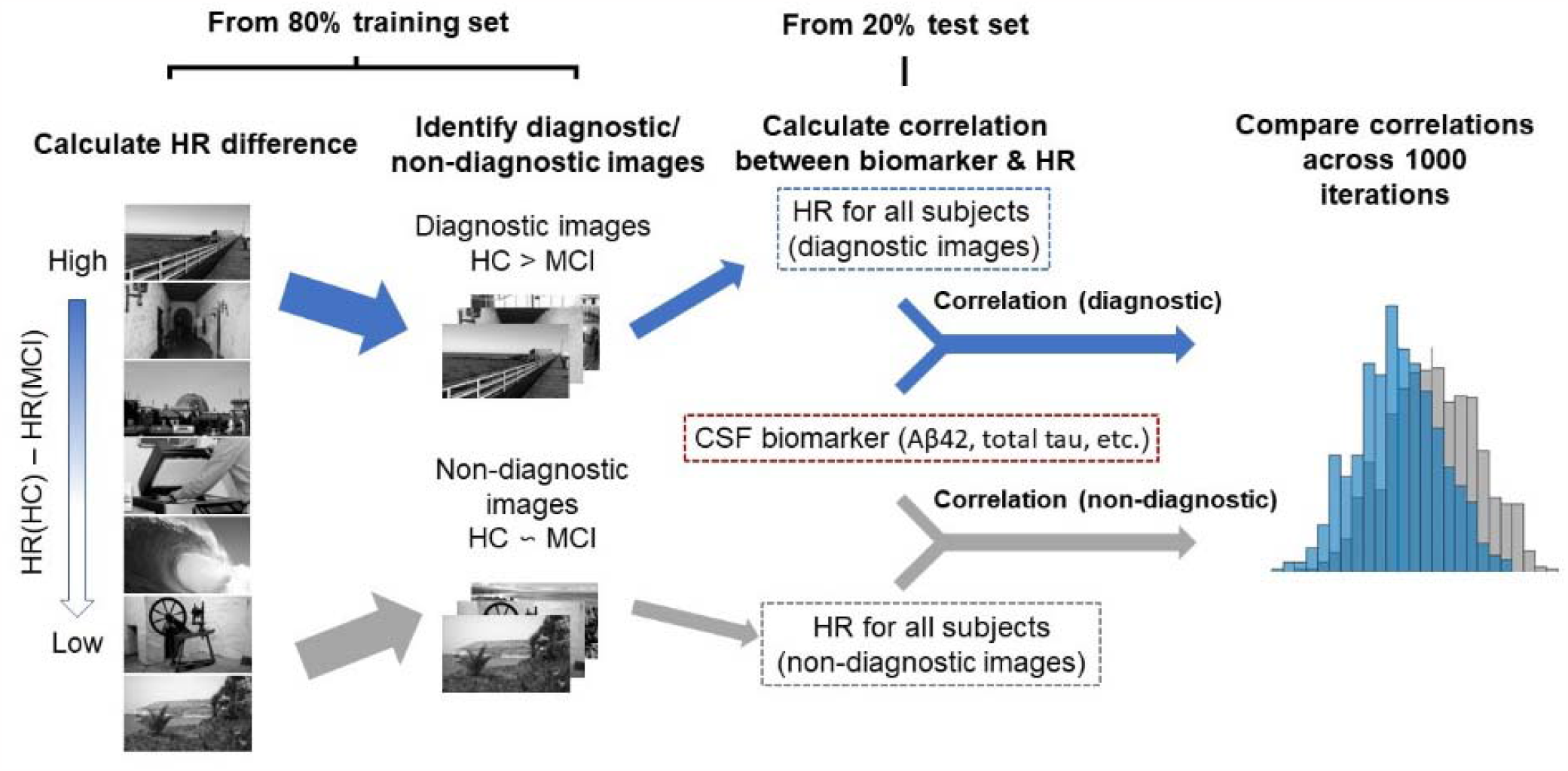
Analysis process for the comparison between behavior-biomarker correlations. In the training set (80% of participants, subsampled to have even group membership), 50 diagnostic images and 50 non-diagnostic images were identified. In the test set (the remaining 20% participants), we compared the HR-biomarker correlation of diagnostic images vs. non-diagnostic images.

**Figure 3.**
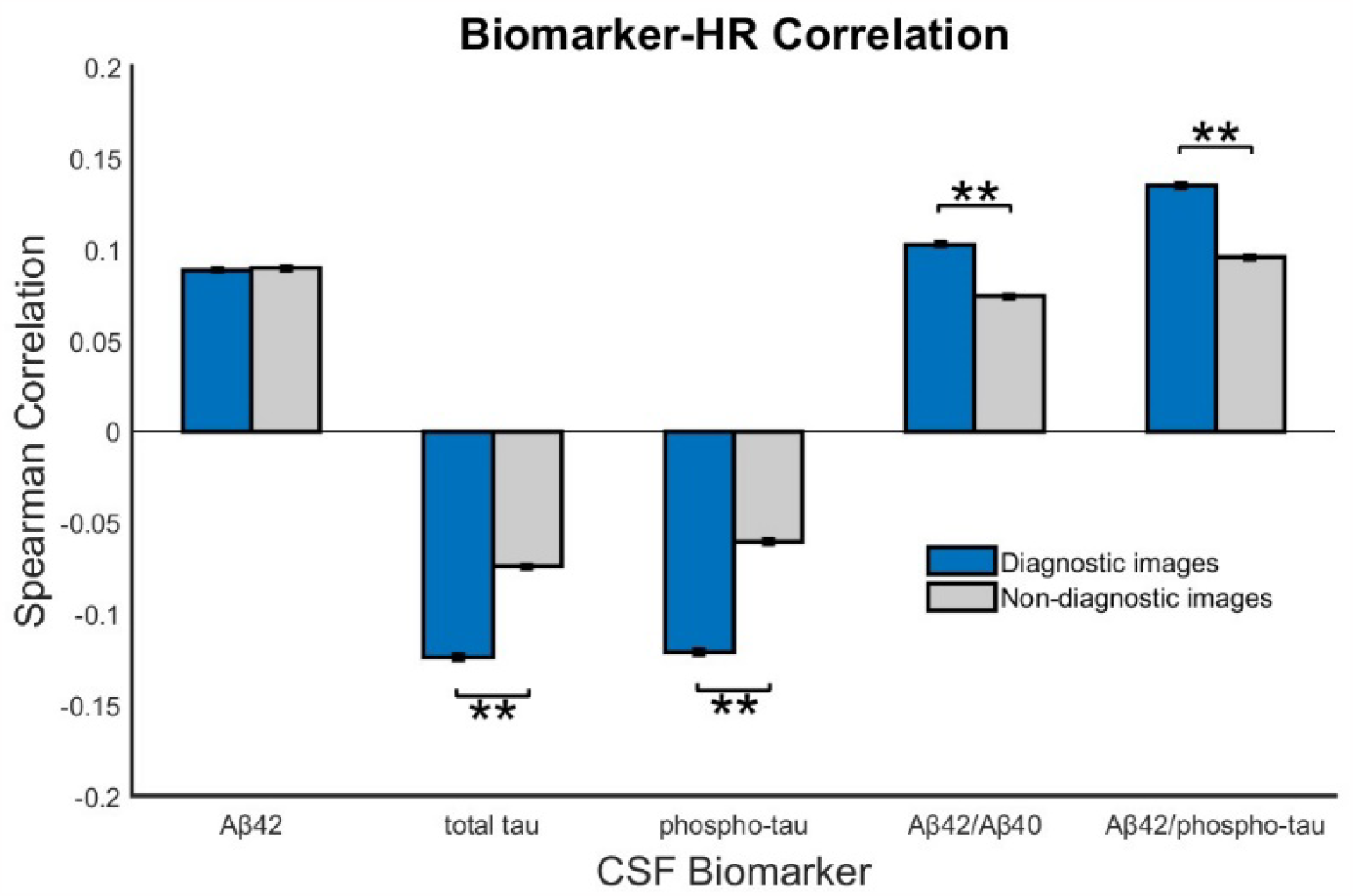
Comparison of biomarker-HR correlation between diagnostic and non-diagnostic images. ** indicate significance at a level of p < 0.001.

### MCI participants have higher activation in left hemisphere scene processing regions for diagnostic images

Through fMRI analysis, we investigated which brain regions contribute to processing diagnostic images. We first conducted a whole-brain univariate analysis to locate potential areas that are related to image diagnosticity. For the HC and MCI groups separately, we averaged the volumes for diagnostic and non-diagnostic images for each individual and conducted a group-level t-test between diagnostic and non-diagnostic images (Figure 4). For MCI participants, several temporal and occipital regions that comprise the ventral visual stream (Kravitz et al., 2011) and overlap with scene processing regions (Silson et al., 2019) in the left hemisphere showed significantly higher activation for diagnostic images compared to non-diagnostic images. No difference was observed in the hippocampus. In HC participants, these scene processing regions showed no differences between diagnostic and non-diagnostic images. This suggests that while HC participants showed no effect of image diagnosticity in these regions, MCI participants had increased activation for the diagnostic images compared to non-diagnostic images, even though they performed worse on the diagnostic images.

**Figure 4.**
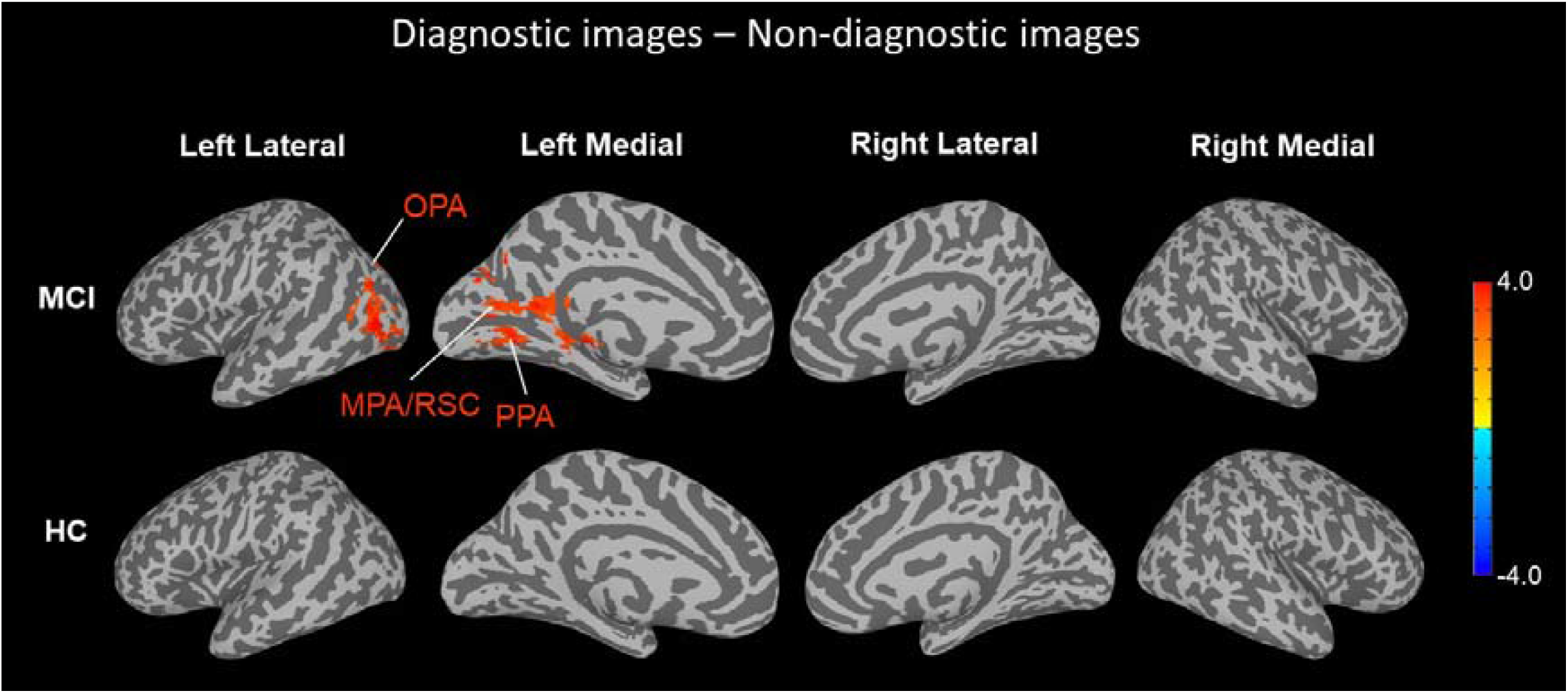
Univariate contrast results (t-statistic) between diagnostic and non-diagnostic images in HC and MCI participants (p < 0.005, cluster threshold corrected at 131 voxels). For MCI participants, diagnostic images showed significantly higher activation in left PPA, MPA/RSC and OPA. HC participants showed no differences in activation between diagnostic and non-diagnostic images. In this whole-brain analysis, there were also no right hemisphere regions that showed a significant difference in either group.

In our stimulus set, the diagnostic and non-diagnostic images were all scene images. Thus, we examined whether this activity increase was present in regions specifically involved in scene processing and memory. We located key scene processing ROIs (PPA, MPA and OPA) in both hemispheres using independent templates (Silson et al., 2019) and conducted a mixed-design ANOVA (participant group * image diagnosticity) within the ROIs (Figure 5). No significant main effect of participant group was found in any of the ROIs, which means that the average activations across all images within scene ROIs were not systematically different between HC and MCI participants. Within the left hemisphere, a significant main effect of image diagnosticity was found in left PPA (F(110) = 5.689, p = 0.019), left MPA (F(110) = 7.32, p = 0.008) and left OPA (F(110) = 4.22, p = 0.042). In the right hemisphere, a main effect of image diagnosticity was marginally significant in right MPA (F(110) = 3.615, p = 0.06), but not significant in the right PPA (F(110) = 0.035, p = 0.85) or right OPA (F(110) = 1.149, p = 0.28). These image diagnosticity effects imply a general trend of higher activation in scene processing for diagnostic images across participant groups. However, importantly we found a significant interaction between participant group (HC or MCI) and image diagnosticity (diagnostic vs. non-diagnostic) in the left PPA (F(110) = 14.509, p = 0.0002), left MPA (F(110) = 6.679, p = 0.001) and left OPA (F(110) = 2.287, p = 0.03). Paired t-tests showed that there was no effect of diagnosticity in the HC group (left PPA: p = 0.23; left MPA: p = 0.92; left OPA: p= 0.91) but a significant effect of diagnosticity within the MCI group (left PPA: p = 0.0007; left MPA: p = 0.003; left OPA: p = 0.01) in left scene ROIs. The interaction effect was of marginal significance in the right OPA (F(110) = 3.729, p = 0.056) and not significant in the right PPA (F(110) = 0.478, p =0.49) or right MPA (F(110) = 1.501, p = 0.22). These results indicate that MCI participants had higher activation in scene processing regions in the left hemisphere for diagnostic images compared to non-diagnostic images. In contrast, HC participants showed no difference of activation for diagnostic and non-diagnostic images in scene processing ROIs.

**Figure 5.**
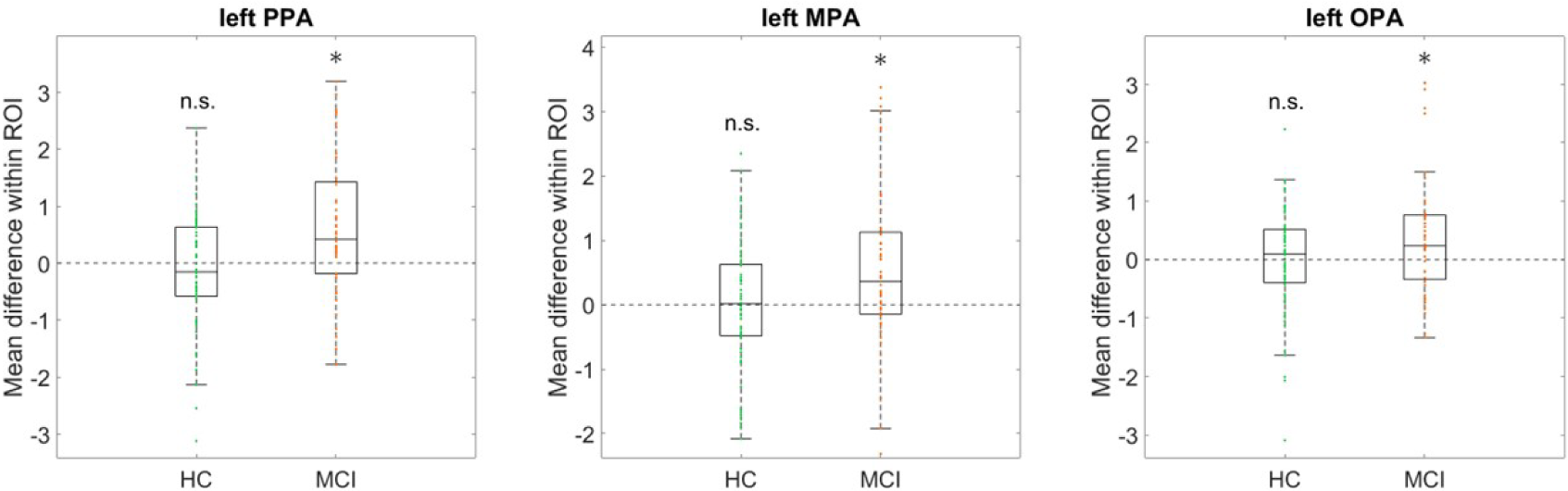
Mean activation difference for diagnostic – non-diagnostic images for HC and MCI participants. In left PPA, MPA/RSC and OPA, MCI participants showed significantly higher activation for diagnostic images compared to non-diagnostic images, while no difference was found in HC participants. * indicates significance at a level of p<0.05.

## Discussion

The aim of this study was to explore whether diagnostic images, which are identified from memorability differences between HCs and MCI participants, are also related to greater sensitivity to CSF biomarker status and differed in brain activation in scene-related regions. We found that the memory performance of diagnostic images had stronger correlations with tau and amyloid beta biomarker status. Among the five CSF biomarkers that we examined, diagnostic images showed significantly higher sensitivity to total tau, phospho-tau, Aβ42/Aβ40 and Aβ42/phospho-tau. These results indicate that both amyloid and tau-pathology contributed to the diagnosticity difference in image memory. We also found that for MCI participants, diagnostic images elicited greater activation than non-diagnostic images in scene-related regions PPA, MPA and OPA in the left hemisphere. In contrast, no difference of brain activation was found in HCs. A hypothesis we propose is that MCI participants have impaired processing of semantically rich scene images and must dedicate more neural resources to processing these images. Thus, MCI participants showed worse memory on these images despite elevated activity in scene-related regions. This hypothesis will need to be validated with further studies directly manipulating the content of scenes to make them more or less diagnostic.

By investigating how memory differs across content, we could potentially get a better understanding of the memory impairment in dementia. Our results show that different aspects of episodic memory (e.g., semantic or perceptual processing) may also have different rates of degradation. Similarly, studies on autobiographical memory have reported a specific impairment of personal thoughts and episodic details compared to semantic details in aging and dementia (Barnabe et al., 2012; Levine et al., 2002; Kopelman et al., 1989). These results also show that differences in memory performance for specific contents are not simply due to the idiosyncratic differences across people or contexts. Instead, this difference is consistent across individuals and can be related to the interplay between stimulus properties and the pathological processes of Alzheimer’s dementia.

In progressive neurodegenerative conditions like Alzheimer’s dementia, the ability to remember certain contents may fade earlier than others, especially when memory ability is not completely lost (Grande et al., 2021). The selective impairment of certain types of stimuli may eventually be attributed to specific sub-components of the memory system. By studying how the memory for certain contents are more affected, we can explore how the architecture of the memory system is affected by pathological aging processes. Moreover, identifying content-specific impairments for certain disease stages will allow better designs for diagnostic assessments. If memory for certain contents is particularly damaged for MCI or Alzheimer’s disease participants, a memory test probing these contents should be more sensitive, as the participants differ more from healthy controls. Our results from the fMRI data revealed greater activations in several scene-related ROIs (MPA, OPA, PPA) for diagnostic images in MCI participants. These data show that it is particularly the scene processing regions that are implicated in memory dysfunction for diagnostic images. Why we observed increased activation in these regions remains to be clarified. Previous studies have reported hyperactivation in hippocampus or parahippocampal gyrus (including PPA) during encoding (Hämäläinen et al., 2006; Kircher et al., 2007) in Alzheimer’s disease. Moreover, MTL hyperactivation is related to the degree of future memory decline in MCI participants (Dickerson & Sperling, 2008). Our analysis is different from these studies as we compared the activation of two sets of images in the same participants. In our analysis, diagnostic images showed increased activation for MCI participants, which may indicate that diagnostic images are those that elicit more hyperactivation than other images even though MCI participants showed difficulties in remembering these images. Thus, increased activation in diagnostic images may be attributed to impairment in scene memory and processing specific to MCI.

For MCI participants, the increased activation of diagnostic images compared to non-diagnostic images appeared mostly in the left hemisphere. While right OPA also showed a marginal effect, most regions in the right hemisphere showed a much smaller effect or no effect. In general, the left hemisphere has been reported to engage in more semantic or verbal processing of memory in healthy participants (Kennepohl et al, 2007; Dalton et al., 2016). For Alzheimer’s disease patients, structural changes and functional connectivity declines seem to be more severe in the left hemisphere. (Berron et al., Liu et al., 2018; Low et al., 2019; Roe et al., 2021). Moreover, Alzheimer’s disease patients with lateralized pathology in the left hemisphere demonstrate more language-specific deficits (Ossenkoppele et al, 2016; Frings et al., 2015). Thus, there might be a link between the level of semantic processing in memory for these specific scene images and the lateralized activation difference between diagnostic and non-diagnostic images. Future research may validate this asymmetry effect and investigate which attributes of the images are most relevant to lateralized brain activity in MCI participants.

There are some limitations to the current work that should motivate future study. First, the measurement of image memorability and diagnosticity scores was noisy due to the limited number of trials assigned to each image. Each participant only saw a limited subset of the whole image pool (88 out of 835 images), resulting in around ten trials per image each for the HC and MCI groups. In this case, the consistency of the memorability score may be lower than the ideal case where each image is viewed by many more participants. Moreover, it is possible that memory performance for MCI participants are less consistent than HCs. It would be important to replicate these findings with an experiment dedicated to a relatively smaller set of images with more participants. It would also be interesting to test how manipulating the experimental context or task around these diagnostic images impacts their ability to diagnose an impairment. Additionally, future study may explore whether semantic processing was the missing component in MCI participants for diagnostic images by testing this relationship in in other image categories (e.g., objects, faces) or directly manipulating the semantic content of images.

Our results show that when HCs and MCI participants take the same memory task, there is a reliable difference in performance driven by the image, which is indicative of neural pathology and reveals differential neural processing. This highlights the importance of measuring and analyzing the diagnosticity of individual images when carrying out memory tests in clinical contexts. If an episodic memory test includes more diagnostic images rather than including random images, it should have higher sensitivity and accuracy. It is also promising to consider content-specific diagnosticity on other clinically oriented tests of memory. More broadly, a content-specific exploration of memory for Alzheimer’s dementia and other dementias (i.e., testing questions of *what* is being remembered and *why*) introduces novel questions and perspectives that are not covered by the conventional view of the memory system. By studying the characteristics of episodic memory in a content-based space, we can explore the functional architecture of the memory system in ways that may have been otherwise impossible to observe.

In conclusion, the current study reveals that diagnostic images for MCI showed a stronger relationship between memory performance and CSF Alzheimer’s disease biomarkers in HC and MCI participants. Moreover, MCI participants have elevated neural activation in scene-related regions for diagnostic images than non-diagnostic images. Our results show that by looking at how memory contents are differentially impaired in MCI, we can better understand the effect of neurodegeneration processes in the memory system.

## Online Methods

### Participants

Behavioral and fMRI data from a visual memory task as well as CSF biomarker data were analyzed from the DZNE Longitudinal Cognitive Impairment and Dementia Study (DELCODE), which is a longitudinal and observational clinical study of Alzheimer’s Disease based in 10 sites in Germany. Details about the study design, data acquisition and tasks were reported in Jessen et al. (2018). We included all healthy control (HC) and mild cognitive impairment (MCI) participants that have both CSF biomarker and fMRI data available, with a total of 64 HC (36 females, 28 males) and 48 MCIs (23 females, 25 males). MCI participants were recruited by referral and self-referral to a memory clinic, while HC participants were recruited by public advertisements. MCI individuals all reported memory concerns and performed more than 1.5 standard deviations below the age-, sex-, and education-adjusted mean in the delayed recall subtest of the Consortium to Establish a Registry for Alzheimer’s Disease (CERAD) neuropsychological battery. HC individuals had normal cognitive performance (within 1.5 SD) in all CERAD-NP subtests and had no subjective concerns of decline in cognitive functioning.

### Visual memory task

Participants engaged in a visual memory task with an encoding phase in the scanner and a recognition phase outside the scanner. In the encoding phase, the participants viewed 88 novel scene images (44 indoor and 44 outdoor) as well as 22 repetitions for each of two pre-familiarized scene images in random order. We do not report any analyses on the pre-familiarized scene images, only focusing on novel images. Images were 8-bit grayscale indoor (e.g., rooms, halls) or outdoor (e.g., natural scene, buildings) scene images, occasionally with people in the scene. Images were presented on a 30” MRI-compatible LCD screen (Medres OptoStim), scaled to 1250 * 750-pixel resolution and matched for distance, luminance, color and contrast, with a viewing horizontal half-angle of 10.05° across scanners. Each image was displayed for 2500ms with an optimized jitter for statistical efficiency. Participants were asked to categorize images as ‘indoor’ or ‘outdoor’ with button presses. In the recognition phase after a 70-min delay, participants provided recognition ratings for the 88 scene images from the encoding phase and 44 novel foil scene images. The rating was based on a 5-point scale: (1) I am sure that this picture is new, (2) I think that this picture is new, (3) I cannot decide if this picture is new or old, (4) I think I saw this picture before, or (5) I am sure that I did see this picture before. A 1 or 2 rating was coded as a new judgement, while a 4 or 5 rating was coded as an old judgement. All images used in the encoding phase and the recognition phase (including foil images) were randomly selected for each participant from an image pool with a total of 835 images.

### CSF Alzheimer’s disease biomarker assessment

CSF Alzheimer’s disease biomarkers were collected in the DELCODE dataset for the set of participants that we included in our analysis (Jessen et al., 2018). Five types of CSF Alzheimer’s disease biomarkers were determined using commercially available kits according to vendor specifications: V-PLEX Aβ Peptide Panel 1 (6E10) Kit (K15200E) and V-PLEX Human Total Tau Kit (K151LAE) (Mesoscale Diagnostics LLC, Rockville, USA), and Innotest Phospho-Tau (181P) (81581; Fujirebio Germany GmbH, Hannover, Germany). These biomarkers address amyloid pathology (Aβ42, Aβ42/ Aβ40, Aβ42/phospho-tau) or tau pathology (total tau and phospho-tau), which are two hallmarks of the neuronal degeneration process in Alzheimer’s dementia.

### Selecting diagnostic and non-diagnostic images

Using a similar approach to Bainbridge et al., 2019, we identified a diagnostic image subset and a non-diagnostic image subset from the 835-image pool based on the memorability differences between HC and MCI participants. In the biomarker analysis, we selected images based on 80% participants in the training set of each iteration, while in the fMRI analysis we identified the images based on the behavior from all participants. First, hit rate (HR) was calculated within the HC and MCI groups for each image. Images were then sorted by the difference in HR between the HC and MCI groups. Only images with at least three responses from each group were included. The diagnostic subset includes the images for which HC have a much higher hit rate than MCI participants. In other words, HC participants have much better memory performance than MCI participants for these images, compared to any other images in the pool. The non-diagnostic subset includes the images that are in the middle of the sorted sequence. For these ‘median’ images, HC only have a slightly better memory performance than MCI participants, which corresponds to the fact that on average HC have higher memory performance than MCI participants. Bainbridge et al. (2019) found that the diagnostic images by this same definition were better at categorizing HC and MCI participants in receiver operating characteristic analysis of the behavioral data, showing primary evidence that these images may better indicate Alzheimer’s disease-related impairment in cognitive processing.

### Split-half consistency analysis

To examine whether image memorability and diagnosticity are consistent across participants and intrinsic to the images, we conducted a split-half consistency analysis for memorability and diagnosticity within the HC and MCI groups. The analysis consists of 1000 iterations. In each iteration, subjects were randomly split into two halves while keeping the number of HC and MCI participants in each half equal to the other half. For each participant half, we calculated for each image the proportion of correct recognition responses in HCs (HR within HC group) and in MCI participants (HR within MCI group), and the HR difference between the HC and MCI groups. We calculated Spearman rank correlations for each score (HR for HC, HR for MCI, HR difference) between the two participant halves per iteration. Null distributions were also generated by randomly shuffling the scores for one participant half for each score. The correlations between the two halves were Fisher Z-transformed and compared with the null distribution using independent samples t-tests. If the correlation of HR or HR difference between the two random halves have a significant positive correlation, it would indicate that certain images are reliably more memorable / diagnostic across all participants in the sample. This would show that the memory difference between images can be attributed to images themselves rather than mere noise.

### Comparing behavior-biomarker correlations between diagnostic and non-diagnostic images

In this analysis, we compared the magnitude of the correlation between image memory and biomarker measures, separately for diagnostic and non-diagnostic images for the same set of participants. Figure 2 demonstrates the major steps of this biomarker analysis. With a hold-out cross validation approach, we assigned 80% participants to the training set and 20% participants to the test set with a sampling process that enables an equal ratio of HC to MCI participants in the two sets. We identified subsets of 50 diagnostic and 50 non-diagnostic images with the behavioral data of the training set. We then compared the behavior-biomarker correlation of the two sets of images in the test set. Individual participant memory performance was measured by HR within the diagnostic and non-diagnostic subsets separately. We then calculated Spearman correlations between the participants’ HRs and each of the five CSF biomarkers (Aβ42, total tau, phospho-tau, Aβ42/Aβ40, and Aβ42/phospho-tau). We repeated this process 1000 times and generated correlation coefficients for diagnostic and non-diagnostic images for each iteration. Meanwhile, we also calculated the correlation between diagnostic/non-diagnostic image performance and randomly shuffled biomarker scores to generate null distributions. To examine whether the behavior-biomarker correlation for the diagnostic set is significantly higher than that for non-diagnostic images, we subtracted the Fisher Z-transformed correlation coefficients of diagnostic and non-diagnostic images and got 1000 correlation score differences for each biomarker. Similarly, a null distribution was also generated from shuffled data. This distribution of correlation differences was compared with a null distribution using an independent samples t-test.

### MRI acquisition and preprocessing

MRI data were acquired at nine scanning sites across Germany with Siemens scanners (three TIM Trio systems, four Verio systems, one Skyra and one Prisma system). A 5-minute long T1-weighted anatomical image was acquired with the following specifications: 3D GRAPPA PAT 2, 1 mm^3^ isotropic voxels, 256 × 256 px, 192 slices, sagittal, repetition time (TR) 2500 ms, echo time (TE) 4.33 ms, inversion time (TI) 110 ms, flip angle (FA) 7°. A 12-minute long T2-weighted anatomical image was also acquired: optimized for medial temporal lobe volumetry, 0.5 × 0.5 × 1.5 mm_3_ voxels, 384 × 384 px, 64 slices, orthogonal to the hippocampal long axis, TR 3500 ms, TE 353 ms. Participants engaged in one run of task-based functional MRI scan including a scene novelty and encoding task: 2D echo planar imaging (EPI), GRAPPA PAT 2, 3.5 mm^3^ isotropic voxels, 64 × 64 px, 47 slices, oblique axial/AC-PC aligned, TR 2580 ms, TE 30 ms, FA 80°, 206 volumes. See Jessen et al. (2018) and Düzel et al. (2018) for more details about these fMRI data.

During the functional scans, participants performed a task where they viewed 132 scene images (88 target images, 44 repetition images) and were instructed to make indoor/outdoor judgement for each image (described above in “Visual Memory Task”). All scanning sites used the same 301111 MR-compatible LCD screen (Medres Optostim) matched for distance, luminance, color and contrast across sites, and the same response buttons (CurrentDesign). All participants underwent vision correction with MR-compatible goggles (MediGlasses, Cambridge Research Systems) according to the same standard operating procedures (SOP). SOPs, quality assurance and assessment (QA) were provided and supervised by the DZNE imaging network (iNET, Magdeburg) as described in Jessen et al. (2014).

### Whole brain univariate analyses

To explore the brain activation differences between diagnostic and non-diagnostic images, we first compared the fMRI data of these two sets of images within the HC and MCI groups across the whole brain. Specifically, we conducted group-level univariate analyses for diagnostic vs. non-diagnostic images with MATLAB and AFNI. We first identified 100 diagnostic and 100 non-diagnostic images from the entire pool of 835 images based on the behavioral performance from the whole dataset and labeled the trials that contained either these diagnostic or non-diagnostic images for each participant. On average, each participant viewed 9 diagnostic or non-diagnostic images and each diagnostic or non-diagnostic image was seen by 7 HC participants and 5 MCI participants. At the individual subject level, we averaged all brain volumes for diagnostic and non-diagnostic images separately. The averaged volumes were then co-registered to the Montreal Neurological Institute (MNI) 2009 template using participants’ corresponding T1 images. After that, we conducted a paired t-test between participants’ diagnostic and non-diagnostic volumes to draw group-level inferences in the HC and MCI groups. Whole brain maps were thresholded at a level of α=0.005 and corrected for multiple comparisons with a cluster threshold corrected cluster size of 131 voxels.

### ROI analyses

Besides an exploratory contrast analysis across the brain, we are specifically interested in how image diagnosticity affects visual processing and memory in the brain. Given the stimuli were all scene images, we analyzed three regions of interest (ROIs) related to scene perception and memory: the PPA (parahippocampal place area), RSC/MPA (retrosplenial complex / medial place area) and TOS/OPA (transverse occipital sulcus / occipital place area). These regions respond selectively to scenes with functional distinctions across them (Epstein & Baker, 2019). The PPA is thought to be involved in scene perception and navigation, the MPA is involved in scene perception, navigation and scene memory, and the OPA is mainly related to perceptual scene processing. ROI voxel specification data (demarcating the three ROIs in the left and right hemispheres) were extracted from Silson et al. (2019) and transformed into the MNI space. For each participant, the average activations for diagnostic and non-diagnostic images were calculated within each ROI. A mixed-design ANOVA (within-group: diagnostic/non-diagnostic images; between group: HC/MCI participant) was used to explore the effect of image diagnosticity as well as the interaction effect on ROI activation for each of the left and right ROIs.

## Acknowledgements

Many thanks to Monica D. Rosenberg for her insightful feedback on this work, and to Hartmut Schuetze on his support with data access and management.

## References

Bainbridge, W.A. (2020). The resiliency of image memorability: A predictor of memory separate from attention and priming. Neuropsychologia, 141, 107408.

Bainbridge, W.A., Berron, D., Schütze, H., Cardenas-Blanco, A., Metzger, C., Dobisch, L., et al. (2019). Memorability of photographs in subjective cognitive decline and mild cognitive impairment for cognitive assessment. Alzheimer’s & Dementia: Diagnosis, Assessment & Disease Monitoring, 11, 610–618.

Bainbridge, W.A., Dilks, D.D., & Oliva, A. (2017). Memorability: A stimulus-driven perceptual neural signature distinctive from memory. NeuroImage, 149, 141–152.

Bainbridge, W.A., Isola, P., & Oliva, A. (2013). The intrinsic memorability of face photographs. Journal of Experimental Psychology: General, 142, 1323–1334.

Bainbridge, W.A., & Rissman, J. (2018). Dissociating neural markers of stimulus memorability and subjective recognition during episodic retrieval. Scientific Reports, 8, 8679.

Barnabe, A., Whitehead, V., Pilon, R., Arsenault-Lapierre, G., Chertkow, H., 2012. Autobiographical memory in mild cognitive impairment and Alzheimer’s disease: a comparison between the Levine and Kopelman interview methodologies. Hippocampus 22, 1809–1825. 10.1002/hipo.22015.

Berron, D., Cardenas-Blanco, A., Bittner, D., Metzger, C.D., Spottke, A., Heneka, M.T., Fliessbach, K., Schneider, A., Teipel, S.J., Wagner, M., Speck, O., Jessen, F., Düzel, E., 2019. Higher CSF tau levels are related to hippocampal hyperactivity and object mnemonic discrimination in older adults. J. Neurosci. 39, 8788–8797. 10.1523/JNEUROSCI.1279-19.2019.

Berron, D., van Westen, D., Ossenkoppele, R., Strandberg, O., & Hansson, O. (2020). Medial temporal lobe connectivity and its associations with cognition in early Alzheimer’s disease. Brain, 143(4), 1233–1248.

Borkin, M. A., Bylinskii, Z., Kim, N. W., Bainbridge, C. M., Yen C. S., Borkin, D., Pfister, H., & Oliva, A. (2015). Beyond memorability: Visualization recognition and recall. IEEE Transactions on Visualization and Computer Graphics, 22, 519–528.

Braak, H., Alafuzoff, I., Arzberger, T., Kretzschmar, H., Del Tredici, K., 2006. Staging of Alzheimer disease-associated neurofibrillary pathology using paraffin sections and immunocytochemistry. Acta Neuropathol. 112, 389–404. 10.1007/s00401-006-0127-z.

Broers, N., & Busch, N.A. (2020). The effect of intrinsic image memorability on recollection and familiarity. Memory & Cognition, 1–21.

Costa, A., Bak, T., Caffarra, P., Caltagirone, C., Ceccaldi, M., Collette, F., Crutch, S., Della Sala, S., Démonet, J.F., Dubois, B., Duzel, E., Nestor, P., Papageorgiou, S.G., Salmon, E., Sikkes, S., Tiraboschi, P., van der Flier, W.M., Visser, P.J., Cappa, S.F., 2017. The need for harmonisation and innovation of neuropsychological assessment in neurodegenerative dementias in Europe: consensus document of the Joint Program for Neurodegenerative Diseases Working Group. Alzheimer’s Res. Ther. 9, 27. 10.1186/s13195-017-0254-x.

Dalton, M. A., Hornberger, M., & Piguet, O. (2016). Material specific lateralization of medial temporal lobe function: an f MRI investigation. Human brain mapping, 37(3), 933–941.

Dickerson, B. C., & Sperling, R. A. (2008). Functional abnormalities of the medial temporal lobe memory system in mild cognitive impairment and Alzheimer’s disease: insights from functional MRI studies. Neuropsychologia, 46(6), 1624–1635.

Dubey, R., Peterson, J., Khosla, A., Yang, M.H., & Ghanem, B. (2015). What makes an object memorable? Proceedings of the IEEE International Conference on Computer vision, 1089–1097.

Düzel, E., Berron, D., Schütze, H., Cardenas-Blanco, A., Metzger, C., Betts, M., & Jessen, F. (2018). CSF total tau levels are associated with hippocampal novelty irrespective of hippocampal volume. Alzheimer’s & Dementia: Diagnosis, Assessment & Disease Monitoring, 10(1), 782–790.

Frings, L., Hellwig, S., Spehl, T. S., Bormann, T., Buchert, R., Vach, W., & Meyer, P. T. (2015). Asymmetries of amyloid-β burden and neuronal dysfunction are positively correlated in Alzheimer’s disease. Brain, 138(10), 3089–3099.

Goetschalckx, L., Moors, P., & Wagemans, J. (2017). Image memorability across longer time intervals. Memory, 26, 581–588.

Grande, X., Berron, D., Maass, A., Bainbridge, W. A., & Düzel, E. (2021). Content-specific vulnerability of recent episodic memories in Alzheimer’s disease. Neuropsychologia, 160, 107976.

Grothe, M.J., Barthel, H., Sepulcre, J., Dyrba, M., Sabri, O., Teipel, S.J., Initiative, F. the A.D.N., 2017. In vivo staging of regional amyloid deposition. Neurology 89, 2031–2038. 10.1212/WNL.0000000000004643.

Hämäläinen, A., Pihlajamäki, M., Tanila, H., Hänninen, T., Niskanen, E., Tervo, S., & Soininen, H. (2007). Increased fMRI responses during encoding in mild cognitive impairment. Neurobiology of aging, 28(12), 1889–1903.

Hansson, O., Lehmann, S., Otto, M., Zetterberg, H., & Lewczuk, P. (2019). Advantages and disadvantages of the use of the CSF Amyloid β (Aβ) 42/40 ratio in the diagnosis of Alzheimer’s Disease. Alzheimer’s research & therapy, 11, 1–15.

Isola, P., Xiao, J., Parikh, D., Torralba, A., & Oliva, A. (2013). What makes a photograph memorable? IEEE Transactions on Pattern Analysis and Machine Learning, 36, 1469–1482.

Isola, P., Xiao, J. X., Torralba, A., & Oliva, A. (2011a). What makes an image memorable? 24th IEEE Conference on Computer Vision and Pattern Recognition (CVPR), 145–152.

Jagust, W., 2018. Imaging the evolution and pathophysiology of Alzheimer disease. Nat. Rev. Neurosci. 19, 687–700. 10.1038/s41583-018-0067-3.

Jessen, F., Spottke, A., Boecker, H., Brosseron, F., Buerger, K., Catak, C., & Düzel, E. (2018). Design and first baseline data of the DZNE multicenter observational study on predementia Alzheimer’s disease (DELCODE). Alzheimer’s research & therapy, 10(1), 1–10.

Jessen, F., Amariglio, R. E., Van Boxtel, M., Breteler, M., Ceccaldi, M., Chételat, G., & Subjective Cognitive Decline Initiative. (2014). A conceptual framework for research on subjective cognitive decline in preclinical Alzheimer’s disease. Alzheimer’s & dementia, 10(6), 844–852.

Kennepohl, S., Sziklas, V., Garver, K. E., Wagner, D. D., & Jones-Gotman, M. (2007). Memory and the medial temporal lobe: hemispheric specialization reconsidered. Neuroimage, 36(3), 969–978.

Kircher, T. T., Weis, S., Freymann, K., Erb, M., Jessen, F., Grodd, W., & Leube, D. T. (2007). Hippocampal activation in patients with mild cognitive impairment is necessary for successful memory encoding. Journal of Neurology, Neurosurgery & Psychiatry, 78(8), 812–818.

Kopelman, M.D., Wilson, B.A., Baddeley, A.D., 1989. The autobiographical memory interview: a new assessment of autobiographical and personal semantic memory in amnesic patients. J. Clin. Exp. Neuropsychol. 11, 724–744. 10.1080/01688638908400928.

Kramer, M. A., Hebart, M. N., Baker, C. I., & Bainbridge, W. A. (2023). The features underlying the memorability of objects. Science advances, 9(17), eadd2981.

Kravitz, D. J., Saleem, K. S., Baker, C. I., & Mishkin, M. (2011). A new neural framework for visuospatial processing. Nature Reviews Neuroscience, 12(4), 217–230.

Lee, A.C.H., Buckley, M.J., Pegman, S.J., Spiers, H., Scahill, V.L., Gaffan, D., Bussey, T.J., Davies, R.R., Kapur, N., Hodges, J.R., Graham, K.S., 2005. Specialization in the medial temporal lobe for processing of objects and scenes. Hippocampus 15, 782–797. 10.1002/hipo.20101.

Levine, B., Svoboda, E., Hay, J.F., Winocur, G., Moscovitch, M., 2002. Aging andautobiographical memory: dissociating episodic from semantic retrieval. Psychol. Aging 17, 677–689. 10.1037/0882-7974.17.4.677.

Liu, H., Zhang, L., Xi, Q., Zhao, X., Wang, F., Wang, X., & Lin, Q. (2018). Changes in brain lateralization in patients with mild cognitive impairment and Alzheimer’s disease: a resting-state functional magnetic resonance study from Alzheimer’s disease neuroimaging initiative. Frontiers in neurology, 9, 3.

Low, A., Mak, E., Malpetti, M., Chouliaras, L., Nicastro, N., Su, L., & O’Brien, J. T. (2019). Asymmetrical atrophy of thalamic subnuclei in Alzheimer’s disease and amyloid-positive mild cognitive impairment is associated with key clinical features. Alzheimer’s & Dementia: Diagnosis, Assessment & Disease Monitoring, 11(1), 690–699.

Maass, A., Berron, D., Harrison, T.M., Adams, J.N., La Joie, R., Baker, S., Mellinger, T., Bell, R.K., Swinnerton, K., Inglis, B., Rabinovici, G.D., Düzel, E., Jagust, W.J., 2019. Alzheimer’s pathology targets distinct memory networks in the ageing brain. Brain 142, 2492–2509. 10.1093/brain/awz154.

Maass, A., Landau, S., Baker, S.L., Horng, A., Lockhart, S.N., La Joie, R., Rabinovici, G.D., Jagust, W.J., Alzheimer’s Disease Neuroimaging Initiative, for the A.D.N, 2017. Comparison of multiple tau-PET measures as biomarkers in aging and Alzheimer’s disease. Neuroimage 157, 448–463. 10.1016/j.neuroimage.2017.05.058.

Mohsenzadeh, Y., Mullin, C., Oliva, A., & Pantazis, D. (2019). The perceptual neural trace of memorable unseen scenes. Scientific Reports, 9, 1–10.

Ossenkoppele, R., Schonhaut, D. R., Schöll, M., Lockhart, S. N., Ayakta, N., Baker, S. L., & Rabinovici, G. D. (2016). Tau PET patterns mirror clinical and neuroanatomical variability in Alzheimer’s disease. Brain, 139(5), 1551–1567.

Roe, J. M., Vidal-Piñeiro, D., Sørensen, Ø., Brandmaier, A. M., Düzel, S., Gonzalez, H. A., & Westerhausen, R. (2021). Asymmetric thinning of the cerebral cortex across the adult lifespan is accelerated in Alzheimer’s disease. Nature communications, 12(1), 721.

Wakeland-Hart, C.D., Cao, S. deBettencourt, M.T., Bainbridge, W.A., & Rosenberg, M.D. (2022). Predicting visual memory across images and within individuals. Cognition, 227, 105201.

Xie, W., Bainbridge, W.A., Inati, S.K., Baker, C.I., & Zaghloul, K. (2020). Memorability of words in arbitrary verbal associations modulates memory retrieval in the anterior temporal lobe. Nature Human Behaviour, 4, 937–948.

